# Estimating the fitness effect of deleterious mutations during the two phases of the life cycle: a new method applied to the root-rot fungus *Heterobasidion parviporum*

**DOI:** 10.1101/492686

**Authors:** Pierre-Henri Clergeot, Nicolas O. Rode, Sylvain Glémin, Mikael Brandström-Durling, Katarina Ihrmark, Åke Olson

**Author notes:** These authors contributed equally to this work and are listed alphabetically (Pierre-Henri Clergeot, Nicolas O. Rode). Corresponding author: CBGP, 755 avenue du Campus Agropolis, CS 30016, 34988, Montferrier sur Lez, France.

## Abstract

Many eukaryote species including taxa such as fungi or algae have a lifecycle with substantial haploid and diploid phases. A recent theoretical model predicts that such haploid-diploid lifecycles are stable over long evolutionary time scales when segregating deleterious mutations have stronger effects in homozygous diploids than in haploids and when they are partially recessive in heterozygous diploids. The model predicts that effective dominance, a measure that accounts for these two effects, should be close to 0.5 in these species. It also predicts that diploids should have higher fitness than haploids on average. However, an appropriate statistical framework to conjointly investigate these predictions is currently lacking. In this study, we derive a new quantitative genetic model to test these predictions using fitness data of two haploid parents and their diploid offspring and genome-wide genetic distance between haploid parents. We apply this model to the root-rot basidiomycete fungus *Heterobasidion parviporum*, a species where the heterokaryotic (equivalent to the diploid) phase is longer than the homokaryotic (haploid) phase. We measured two fitness-related traits (mycelium growth rate and the ability to degrade wood) in both homokaryons and heterokaryons and we used whole-genome sequencing to estimate nuclear genetic distance between parents. Possibly due to a lack of power, we did not find that deleterious mutations were recessive or more deleterious when expressed during the heterokaryotic phase. Using this model to compare effective dominance among haploid-diploid species where the relative importance of the two phases varies should help better understand the evolution of haploid-diploid life cycles.

**Article summary for Issue Highlights:** Many eukaryote species including taxa such as fungi or algae spend a large portion of their life cycle as haploids and as diploids. Clergeot, Rode *et al.* derive a statistical model to test whether deleterious mutations have stronger effects in homozygous diploids than in haploids, whether they are partially recessive in heterozygous diploids and whether diploids have higher fitness than haploids on average. As an illustration, they use their model to study growth rate and the ability to degrade wood in the root-rot fungus *Heterobasidion parviporum*. Their model should help gaining further insights into the evolution of haploid-diploid life cycles.

## Introduction

The life cycle of sexual eukaryotes is defined by the alternation of haploid and diploid phases. Several eukaryotic taxa, including many fungi and algae species, spend a significant portion of their life cycle both as haploids and diploids (i.e. haploid-diploid life cycle). A recent theoretical model shows that haploid-diploid life cycles can only occur under a restricted set of conditions (Scott and Rescan 2017): i) Haploid-diploid life cycles are stable only when diploids have higher intrinsic fitness compared to haploids (hereafter ploidy effects on fitness). Such fitness differences between haploids and diploids can be due to the fixation of mutations with different fitness effects in haploids vs. diploids due to differences in physiology (Simchen and Jinks 1964; Szafraniec *et al.* 2003; McBride *et al.* 2008; Gerstein 2013; Zörgö *et al.* 2013), in physical characteristics (e.g. cell morphology, Mable 2001), in gene expression (Coelho *et al.* 2007; Von Dassow *et al.* 2009; Rokitta *et al.* 2011; Meng *et al.* 2013; Liu *et al.* 2017) or in ecology (Thornber 2006; Rescan *et al.* 2016). ii) Haploid-diploid life cycles can be stable only when the effect of intrinsic fitness differences, which favors the diploid phase, balances the effect of selection against deleterious mutations, which favors the haploid phase (Figure 1). When few deleterious mutations segregate (i.e. when the haploid phase is long), the higher intrinsic fitness of diploids shifts the balance towards an increased diploid phase. In contrast, when deleterious mutations segregate at high frequencies (i.e. when the diploid phase is long), selection shifts the balance towards an increased haploid phase despite the higher intrinsic fitness of diploids (Figure 1). iii) Assuming that mutations are weakly deleterious (i.e. selection coefficients below 0.1) and that recombination rate is high, haploid-diploid life cycles can be stable only when the average fitness effect of segregating mutations in heterozygous diploids is slightly greater than the average fitness effect of the two haploid parents (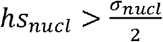, Table 1). Hence, the evolution of haploid-diploid life cycles depend on effective dominance, the product of the average level of dominance of mutations (*h*) and the average of the ratio of fitness effects in homozygous diploids over fitness effects in haploids (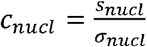, Table 1). Therefore, effective dominance of haploid-diploid species with a high recombination rate can be predicted to be close, but higher than 0.5 (i.e. 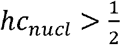), and intrinsic fitness of diploids to be slightly higher than intrinsic fitness of haploids. For example, if mutation are co-dominants 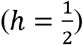, we expect mutations to be more deleterious in homozygous diploids than in haploids 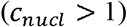.

**Figure 1.**
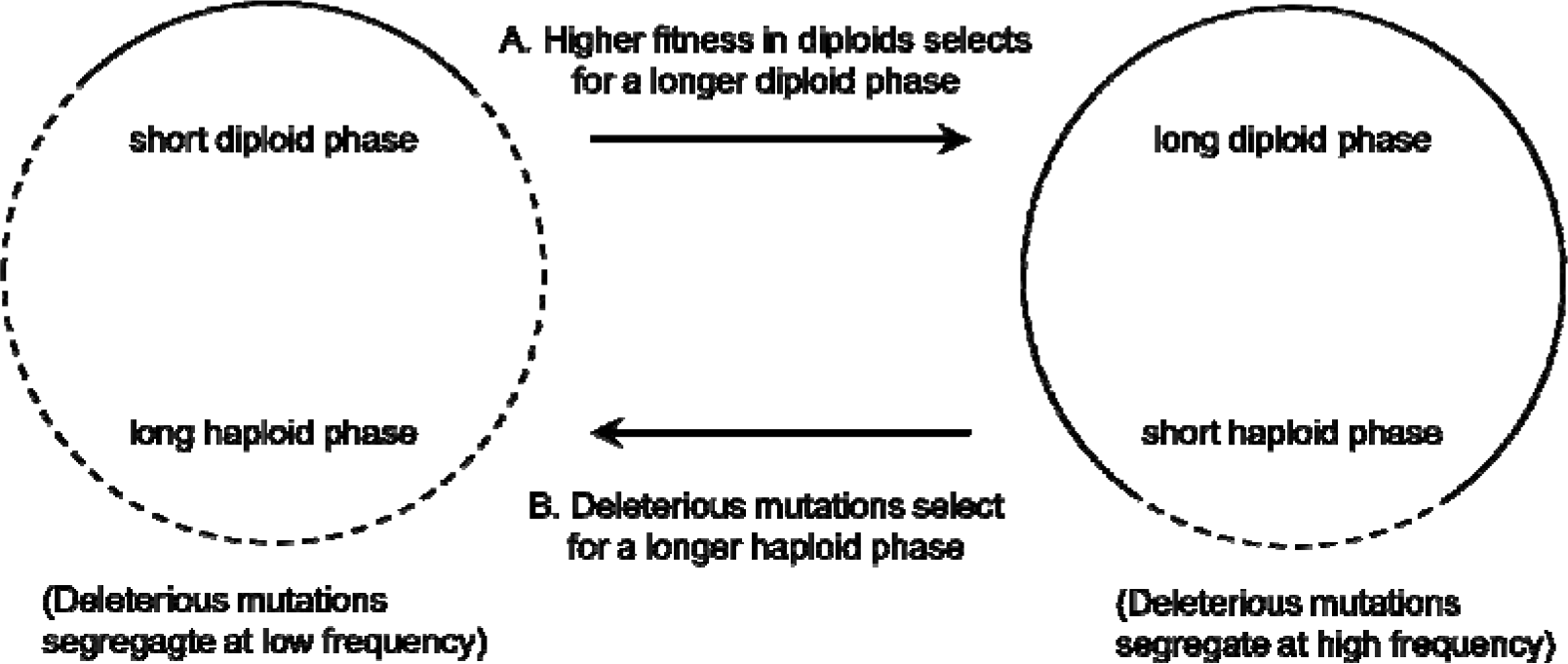
Two selective forces maintain the long-term stability of haploid-diploid life cycles. A. A higher intrinsic fitness in diploids selects for an increase of the diploid phase of the life cycle B. Selection against deleterious mutations favors an increase of the haploid phase. Note that stability requires the effect of mutations in heterozygotes diploids to be slightly higher than the average fitness of the two haploid parents when recombination rate is high (i.e. effective dominance should be slightly higher than 0.5). Stability also requires diploids to have a slightly higher intrinsic fitness than haploids. When diploids have lower intrinsic fitness than haploids, the cycle evolves either towards haplonty or diplonty (see Scott and Rescan 2017 for details).

**Table 1.** Fitness of haploid and diploid genotypes at a nuclear locus and at a mitochondrial locus under selection.

|  | GENOME | GENOTYPE | FITNESS |
| --- | --- | --- | --- |
| **HAPLOID**<br>**SELECTION** | Nucleus | A | $w_A = 1$ |
| | Nucleus | a | $w_a = 1 - \sigma_{nucl}$ |
| | Mitochondrion | $A'$ | $w_{A'} = 1$ |
| | Mitochondrion | $a'$ | $w_{a'} = 1 - \sigma_{mit}$ |
| **DIPLOID**<br>**SELECTION** | Nucleus | AA | $w_{AA} = 1$ |
| | Nucleus | Aa | $w_{Aa} = 1 - h s_{nucl}$ |
| | Nucleus | aa | $w_{aa} = 1 - s_{nucl}$ |
| | Mitochondrion | $A'$ | $w_{A'} = 1$ |
| | Mitochondrion | $a'$ | $w_{a'} = 1 - s_{mit}$ |

$\sigma_{nucl}$ and $s_{nucl}$ : fitness effects of a nuclear mutation in haploids and homozygous diploids. $\sigma_{mit}$ and $s_{mit}$ : fitness effects of a mitochondrial mutation in haploids and diploids. $h$ : dominance of nuclear mutations in diploids. Effective dominance is defined as $hc_{nucl}$ with $c_{nucl} = \frac{s_{nucl}}{\sigma_{nucl}}$ and is predicted to be slightly higher than 0.5 in species with high recombination rate ( $hc_{nucl} > \frac{1}{2}$ see main text for details).

Despite the importance of considering both ploidy effects (differences in intrinsic fitness between haploids and diploids) and effective dominance (*hc*_*nucl*_) to explain the maintenance of haploid-diploid life cycles, these different factors have always been considered separately in experimental studies. For example, studies of yeast natural isolates have estimated the average level of dominance of mutations (Zörgö *et al.* 2012; Plech *et al.* 2014; Shapira *et al.* 2014; Stelkens *et al.* 2014; Bernardes *et al.* 2017) separately from ploidy effects (Zörgö *et al.* 2013). To date, effective dominance has never been estimated experimentally (Scott and Rescan 2017).

A comprehensive theoretical statistical framework to estimate ploidy effects on fitness, average levels of dominance and the ratio of mutation effects on fitness in diploids vs. haploids is currently lacking. Developing such a framework is essential for several reasons. First, it would bring reliable estimations of the components of effective dominance (i.e. *h* and *c*_*nucl*_) based on various types of experimental data (mutation accumulation experiments, multiple mutants, crosses between natural isolates). Comparing estimates based on different approaches is important since the fitness effects of mutations considered in mutation accumulation experiments, knock out mutants, and in crosses between natural isolates could vary greatly. Second, such a quantitative genetic framework would account for the relatedness among haploid parents and among their diploid offspring when estimating ploidy effects and dominance levels. For example, studies on levels of dominance usually assume that fitness of different diploid hybrids are independent, even when some of them share one haploid parent (e.g. Korona 1999; Zörgö *et al.* 2012; Plech *et al.* 2014; Shapira *et al.* 2014; Stelkens *et al.* 2014; Bernardes *et al.* 2017). Such non-independence between fitness data could potentially bias estimates of average levels of dominance. Third, such framework would broaden the potential number of taxa available for the quantification of ploidy effects. Indeed, current methods used to investigate ploidy effects rely on the comparison of autozygous offspring (resulting from the fusion of identical haploid nuclei) with heterozygous offspring (resulting from the fusion of different haploid nuclei). As autozygous offspring cannot be recovered in self-incompatible species, most studies of ploidy effects currently suffer from the confounding masking of deleterious mutations in diploids (Scott and Rescan 2017). Finally, analyzing biological data requires to statistically control for important factors not considered in Scott and Rescan’s model (e.g. potential mitochondrial effects and other cytoplasmic effects, segregation of mutations with either haploid-specific or diploid-specific fitness effects, epistasis between mutations, epigenetic effects, etc.). As these factors can potentially bias the estimates of the parameters of interest, it is important to statistically control for them.

In this study, we derive a quantitative genetic model to estimate ploidy effects on fitness, the average level of dominance and the ratio of fitness effects in diploids vs. haploids based on haploid and diploid fitness data. Our statistical model accounts for the genetic relatedness between ploidy levels (haploid parent and diploid offspring) and among the different haploid and diploid strains tested.

We use this model to estimate ploidy effects on fitness and effective dominance (*h* and *c*_*nucl*_) in the basidiomycete filamentous fungus, *Heterobasidion parviporum.* In filamentous fungi, a mycelium with a single type of nucleus is referred to as a homokaryon, whereas mycelium with two different types of haploid nuclei (which results from the fusion between mycelia with different mating types) is referred to as a heterokaryon (Figure 2). Heterokaryons can be considered as functionally equivalent to diploids, although their different haploid nuclei do not fuse, and although the ratio of their two types of nuclei can be different from 1:1, (Beadle and Coonradt 1944; Raper 1966; Day and Roberts 1969). Basidiomycete fungi comprise many self-incompatible species where heterokaryons are predominant in the life cycle (Figure 2, Crockatt *et al.* 2008). In addition, homokaryons and heterokaryons often have the same ecological niche (e.g. Garbelotto *et al.* 1997; Crockatt *et al.* 2008). Species of the genus *Heterobasidion* have the potential to colonize and degrade the wood of host trees both as homokaryons and heterokaryons (Garbelotto *et al.* 1997; Redfern *et al.* 2001), but *H. parviporum*, the causing agent of a severe root and butt rot disease of spruce, is in nature mainly found as heterokaryon (Johannesson and Stenlid 2004). Wind-dispersed homokaryotic basidiospores establish in tree injuries or on stump surfaces, where they persist transiently until they decay or randomly mate to form heterokaryons (Johannesson and Stenlid 2004). Based on theoretical work by Scott and Rescan (2017), effective dominance is predicted to be slightly higher than 0.5 in *H. parviporum,* as heterokaryons are predominant and as the recombination rate is relatively high in *Heterobasidion* spp. (Lind *et al.* 2012).

**Figure 2.**
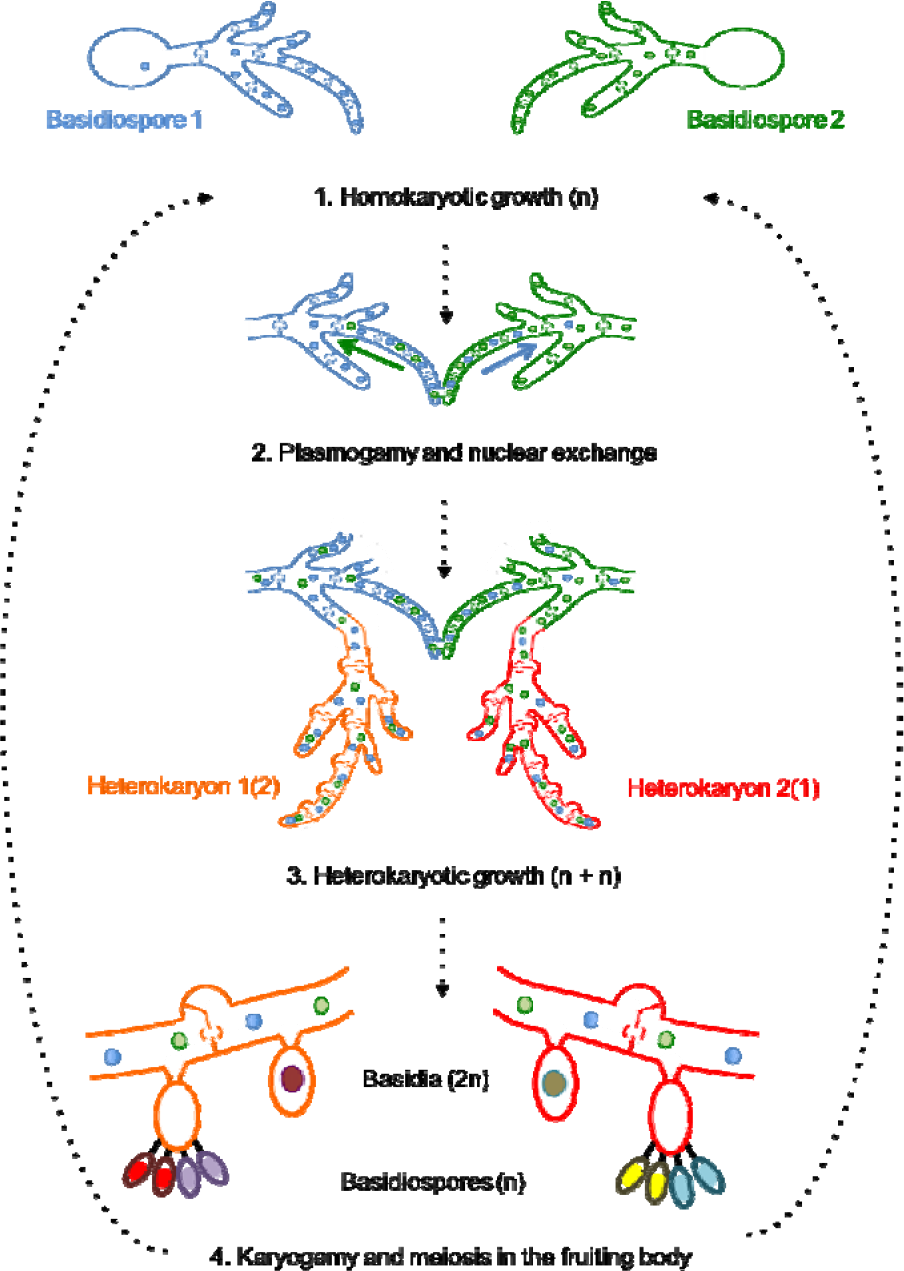
Typical biphasic life cycle of a heterothallic basidiomycete e.g. *Heterobasidion parviporum.* Sexual spores (basidiospores) germinate and form haploid mycelia (depending on the species, these may harbor one nucleus per cell and are named monokaryons, or more than one nucleus, and are named homokaryons). Homokaryons with dissimilar mating types can mate and form a heterokaryotic mycelium by cytoplasmic fusion and reciprocal migration of the haploid nuclei into the opposite parental mycelia. Both nuclear types divide in terminal hyphae during vegetative growth of the heterokaryon, and clamp connections form during cytokinesis to ensure repartition of the two distinct nuclei between mother and daughter cells. Depending on the species, the division of the two different nuclei may be simultaneous (resulting in a nuclear ratio of 1:1) or non-simultaneous (resulting in a nuclear ratio different from 1:1). The former is referred to as dikaryotic, whereas the latter is referred to as heterokaryotic. The heterokaryon eventually develops into a fruiting body, where karyogamy takes place in basidia, the zygotic cells. Meiosis immediately follows karyogamy in basidia, resulting in the formation of new haploid basidiospores.

In both homokaryons and heterokaryons, we measured mycelium growth rate (hereafter MGR), a trait often used as a proxy for fitness in filamentous fungi (Pringle and Taylor 2002). The lower MGR observed in inbred compared to outbred crosses of *Agaricus bisporus* suggests that this trait is related to fitness in basidiomycetes (Xu 1995). We also measured the capacity to degrade spruce wood, a trait usually considered as an important fitness component in *Heterobasidion* spp. (Olson *et al.* 2012). We estimated ploidy effects on fitness, the average level of dominance and the ratio of fitness effects in diploids vs. haploids, and verified the consistency between ecological observations (Johannesson and Stenlid 2004) and predictions based on population genetic theory (Scott and Rescan 2017).

## THEORETICAL QUANTITATIVE GENETIC MODEL

We express the fitness of a diploid genotype as a function of the fitness of its two haploid parents. Details of the derivations are provided in Supplemental Material File S1. We consider that either a wild type or a mutant allele (respectively A and a) is located at each of *N* loci that determine fitness. We assume that the mutant allele at each locus has different fitness effects when present in a haploid (*σ*) or in a homozygous diploid (*s*) individual, but that these fitness effects are constant across the *N* selected loci (Table 1).

Assuming that fitness effects are small and act multiplicatively across loci, the fitness of a haploid genotype *i* with *n*_*i*_ mutant alleles can be defined as:

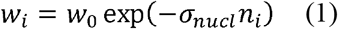

where *σ*_*nucl*_ represent the haploid fitness effect of one mutant alleles at one of the *N* nuclear loci under selection and *w*_0_ represents the baseline fitness of a hypothetical haploid genotype with no deleterious mutations. The effects of mitochondrial mutations can also be added, assuming again small multiplicative effects on fitness:

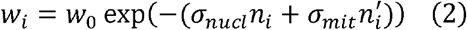

where 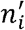 is the number of mitochondrial mutant alleles and*σ*_*mit*_ represent the fitness effect of a mutant alleles at one of the *N*′ mitochondrial loci. Similarly, let *s*_*nucl*_ and *s*_*mit*_ be the homozygote diploid fitness effect of mutant alleles across the *N* nuclear and *N*’ mitochondrial loci. We define the ratio of fitness effects in homozygous diploids vs. haploids for both nuclear 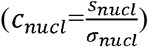 and mitochondrial 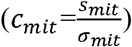 mutations. Hence if *c*_*nucl*_ > 1 or *c*_*mit*_ > 1 (respectively if *c*_*nucl*_ < 1 or *c*_*mit*_ < 1), *s*_*nucl*_ or *s*_*mit*_, the fitness effect of nuclear or mitochondrial deleterious mutant alleles in (homozygous) diploid individuals is larger (respectively smaller) than the fitness effect of nuclear (*σ*_*nucl*_) or mitochondrial (*σ*_*mit*_) deleterious mutant alleles in haploid individuals.

A mating between two haploid genotypes with *n*_*i*_ and *n*_*j*_ nuclear mutant alleles with genotype *i* providing a mitochondrion with 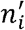 mitochondrial mutant alleles produces a diploid genotype of fitness:

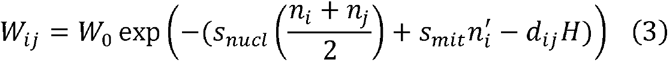

where *W*_0_ represents the baseline fitness of a hypothetical diploid genotype with no deleterious mutations, *s*_*nucl*_ represents the diploid fitness effect of nuclear mutant alleles across the *N* selected loci and *s*_*mit*_ represents the haploid fitness effect of mitochondrial mutant alleles across the *N*′ selected loci, *d*_*ij*_ represents the proportion of selected loci that are heterozygous in the diploid offspring (i.e. the pairwise genetic distance between haploid parents *i* and *j* across the *N* selected loci) and *H* represents a correction term that accounts for dominance 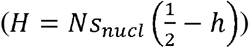. The variable *H*is positive when deleterious mutations are recessive 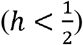 and negative when they are dominant 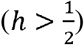. Note that to estimate *h* based on *H,* we need to know both the number of loci selected, *N,* and the average selection coefficient of nuclear mutations, *s*_*nucl*_. To better understand how to interpret *H,* let us take a simple example assuming that a single locus is under selection (*N* = 1). If the first parent has no deleterious mutation (*n*_*i*_ = 0) and the second parent has one deleterious mutation (*n*_*j*_ = 0), we have: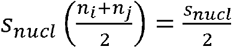 and 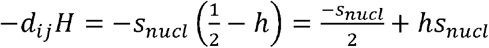. The fitness reduction of the diploid offspring compared to a reference diploid with no deleterious mutation is 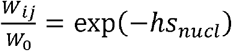. If *hs*_*nucl*_ ≪ 1, this equals 1 – *hs*_*nucl*_ as expected for a heterozygous genotype (Table 1). If we rewrite Eq. (3) using *c*_*nucl*_ and *c*_*mit*_, we have:

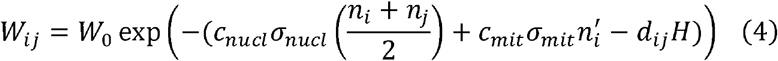

In Supplemental Material File S1, we extend this model to mutations with different fitness effects across the *N* fitness loci. Assuming a large number of loci with small effects, the rationale remains the same but the parameters estimated using Eq. (2) and (4) represent averages of parameters over their respective distributions (i.e. *σ*_*nucl*_ and *σ*_*mit*_ represent the average nuclear and mitochondrial fitness effect in haploids, *H* is positive when mutations are recessive on average and *c*_*nucl*_ is the average across selected loci of the ratio of the fitness effect of each mutation in homozygous diploids over its fitness effects in haploids). We tested the robustness of the predictions of our model when *σ*_*nucl*_, h and *c*_*nucl*_ vary among loci using simulations. The predictions of our model generally hold true in these simulations (see Supplemental File S1-S2 and Figures S1-S2 for details). When the haploid selection coefficients, *σ*_*nucl*_, varies widely among loci, our model tends to slightly overestimate the effect of masking of deleterious mutations in diploid offspring and to slightly underestimate *c*_*nucl*_ (Supplemental File S2, Figure S1B-S2B).

Noting, *A*_*i*_ = -*σ*_*nucl*_*n*_*i*_ and 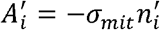 and taking the logarithm, Eq. (2) can then be rewritten as:

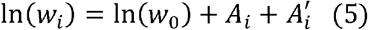

and Eq. (4) can be rewritten as:

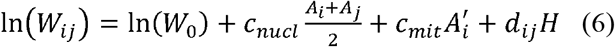

If the number of deleterious nuclear and mitochondrial mutations in homokaryons follows Poisson distributions respectively with means *λ* and *λ*′, the variances among *A*_*i*_ and 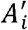 are 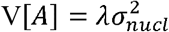 and 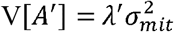 respectively. When the position of deleterious loci is not known and if we assume that fitness is determined by a large number of loci, we can substitute *d*_*ij*_ by *pD*_*ij*_, the genetic distance computed across the whole genome multiplied by a correction factor that is equal to the ratio of average heterozygosity at the *N* selected allele over average heterozygosity over the whole genome (see Supplemental Material File S1 for details). According to Eq. (6) and as the factor *p* is positive, when mutations are recessive on average (H > 0), we expect the fitness of diploid genotypes to increase with the pairwise genetic distance between the two haploid parental genotypes (see Zörgö *et al.* 2012; Plech *et al.* 2014; Shapira *et al.* 2014; Stelkens *et al.* 2014; Bernardes *et al.* 2017 for similar insights based on more verbal predictions). Our model allows testing for intrinsic differences between haploids and diploids (i.e. difference in mean fitness between diploids, *W*_0_, and haploids, *w*_0_) and for different fitness effects of mutations in haploids vs. diploids (i.e. through differences in the variance among haploids vs. the variance among diploids). For exhaustivity, we present a more complex model that accounts for potential epistatic effects between deleterious nuclear mutations (see Supplemental Material File S1 for details). However, in this last model, some parameters are not identifiable using our *H. parviporum* dataset.

## Material and Methods

### Fungal isolates, medium and culture conditions

Thirty homokaryotic isolates of *H. parviporum* sampled at 10 geographic locations in Eurasia over 20 years were used in this study (Table S1). Gene flow between sampling areas is likely, as none of these areas is isolated from the others by large geographic barriers. All samples originated from a single spore colony started from either a basidiospore, or a conidiospore. Homokaryotic and heterokaryotic isolates were routinely maintained on Hagem medium (Stenlid 1985) at 20°C in the dark and stored at 4°C on Hagem medium in glass vials.

### Population structure and genetic distances between homokaryotic isolates

Details regarding the genotyping of the 30 isolates are provided in Supplemental Methods S3. Briefly, in 2012 and 2013, genomic libraries were prepared with DNA from mycelium of each homokaryotic isolate in order to generate 150 bp-long paired-end reads on an Illumina HiSeq platform. In addition, the genome of the reference isolate Sä_159-5 was sequenced using Pacific Biosciences technology, assembled *de novo* using the HGAP3 algorithm (Chin *et al.* 2013), and subsequently corrected by aligning all Sä_159-5 Illumina reads to it. Illumina sequence reads corresponding to each isolate were mapped onto this reference genome using Bowtie2 v2.2.4 (Langmead and Salzberg 2012, see Table S1 for the respective coverage of each isolate). Single nucleotide polymorphisms (SNPs) were called in parallel from the genomes of the 30 isolates using Freebayes v1.0.0-19-gefg685d (Garrison and Marth 2012). SNPs were filtered based on quality, coverage and allelic frequencies (see Supplemental Material S3). Nuclear genetic distance between each pair of isolates was determined using an in-house Perl script as the pairwise sequence divergence calculated over the entire set of filtered SNPs (n > 111,901 nuclear SNPs, Figure S3, see Supplemental Methods S3 for details). Similarly, we computed genetic distance between pairs of mitochondrial haplotypes (n=11 mitochondrial haplotypes, n=145 SNPs, Figure S4) found in the 16 isolates that grew normally on Hagem medium (see below). Finally, we defined four population subgroups corresponding to four large geographic areas (Table S1) and analysed the population structure of this species (see Supplemental Material S3 for details).

### Heterokaryon synthesis and crossing design

Due to self-incompatibility, selfed heterokaryons cannot be recovered in heterothallic fungi. Hence, the 30 homokaryotic isolates were crossed in a pairwise manner according to James et al. (2008), resulting in 870 crosses (Table S4). Briefly, haploid nuclei of homokaryotic mycelia, but not mitochondria, undergo reciprocal migration and exchange during mating so each cross results in two distinct heterokaryons that differ only in their cytoplasm (Ihrmark *et al.* 2002). Hereafter, the homokaryotic isolate that provides only nuclei to a heterokaryon is referred to as heterokaryon donor, whereas the one that provides both nuclei and cytoplasm is referred to as heterokaryon acceptor. Heterokaryon hyphal samples were picked at least two centimeters away from the interaction zone, one month following the initial contact between homokaryotic mycelia. The formation of clamp connections, characteristic of heterokaryotic mycelia, was checked with an optical microscope after sub-cultivation (James et al, 2008). We considered hyphal samples without clamp connections as indicative of the failure of a cross (James et al, 2008). For practical reasons, all heterokaryon syntheses and phenotypic assays were spread over several years (2015-2017).

Overall, 227 of the crosses resulted in the successful formation of heterokaryons that include nuclei from different acceptor and donor parents. Among those, 75 pairs of heterokaryons had identical nuclear background but different mitochondria (i.e. reciprocal crosses where a given strain provided either only nuclei (donor) or both nuclei and mitochondria (acceptor)). Among the 30 homokaryons that grew normally for DNA extractions in 2012/2013, 14 isolates grew very slowly on Hagem medium in 2015-2017 (one of the pre-culture replicates of two isolates also displayed this phenotype, see Table S1). The 14 senescent and 16 non-senescent isolates behaved as nuclei donors for heterokaryon synthesis (Table S1). However, only one senescent and all 16 non-senescent isolates behave as nuclei acceptor (Table S1), this phenomenon being commonly observed in senescent isolates (Stenlid and Rayner 1989; James *et al.* 2008). Mycelium growth rate was measured for the 16 non-senescent homokaryons and 225 heterokaryons (Table S1). The failure to successfully form a heterokaryon might be due to presence of heterokaryon incompatibility alleles or to the senescence phenotype.

Our crossing design is equivalent to unbalanced diallel designs with reciprocals but no selfed crosses, commonly used in quantitative genetics (Lynch and Walsh 1998, p614). The homokaryon genetic effect of a given strain can be defined as the average MGR of homokaryons that carry both the cytoplasm and the nuclei of this strain. The acceptor genetic effect of a given strain can be defined as the average MGR of heterokaryons that carry both the cytoplasm and the nuclei of this strain. Similarly, the donor genetic effect of a given strain is defined as the average MGR of heterokaryons that carry only the nuclei of this strain. Acceptor and donor genetic effects are equivalent to specific combining abilities in diallel breeding designs (Lynch and Walsh 1998, p614), that would be sex-specific. For a genotype, acceptor and donor genetic effects on fitness can indeed differ due to maternal (e.g. mitochondrial or nuclear) effects or to differential nuclear effects in acceptors vs. donors. For example, when the same trait is selected in different directions in donors and acceptors, we expect a negative correlation between acceptor and donor genetic effects (a hallmark of sexual conflicts; Chippindale *et al.* 2001). In addition, due to maternal (e.g. cytoplasmic) effects, one expects acceptor genetic variance to be higher than donor genetic variance (Lynch and Walsh 1998, p601).

### Estimation of mycelium growth rate and wood degradation capacity in homokaryons and heterokaryons

The methods regarding the ability to degrade wood are presented in Supplemental File S2. Confirmed heterokaryons were sub-cultivated twice during one week prior to any experiment to ensure genetic homogeneity of the mycelium (James *et al.* 2008). To estimate the MGR of each of the 16 homokaryons and 225 heterokaryons, we inoculated three Hagem agar plates each with a 4 mm diameter plug taken from the margin of an actively growing pre-culture over different assays (James *et al.* 2008).

For each homokaryon or heterokaryon isolate, we used different precultures for each assay. Plates were incubated at 20°C in the dark. Colony radius was estimated after four, five and six days of growth from four independent measurements per plate. At least two independent assays per isolate were performed (on average, n=2.9 assays per homokaryon isolate and n=2.2 assays per heterokaryon isolate). For each plate and each of the four measurements, growth rates were estimated using a linear regression of the radius over time, which resulted in a total of 552 and 5019 mycelium growth rate estimates for homokaryons and heterokaryons respectively. We used these MGR estimates for statistical analyses (see below). In addition, we estimated the ability of 16 homokaryons and 198 heterokaryons to degrade spruce wood (see Supplemental File S2).

### New statistical model for the estimation of *c*_*nucl*_ *c*_*mit*_, and the average level of dominance of a trait

The theoretical model developed above is general and does not permit the direct inference of the parameters of interest (i.e. *c*_*nucl*_, *c*_*mit*_, the average level of dominance and intrinsic fitness difference between haploids and diploids) based on haploid and diploid fitness data. Indeed, analyzing biological data requires to statistically model important factors that could potentially bias the estimation of these genetic parameters (i.e. mitochondrial effects, variation in fitness due to mutations with homokaryon-specific or heterokaryon-specific fitness effects and the senescence phenotype observed in some homokaryons). To this end, we develop a new custom statistical model to infer the parameters of interest.

Traditionally for diploid organisms, the heritability of a trait relates the phenotype of an offspring and the phenotype of its mid-parent (i.e. the average value of the same trait computed between parents, Lynch and Walsh 1998, p. 8). By analogy, *c*_*nucl*_ and *c*_*mit*_ are related to the heritability between phases of the life cycle. We first develop a model that accounts for nuclear and mitochondrial effects and show that the parameter *c*_*nucl*_ is related to the covariance between the fitness of a heterokaryon and the fitness of its donor homokaryon parent (see Simchen and Jinks 1964 for a seminal study in Schizophyllum commune with a simpler model without mitochondrial effects), while both *c*_*nucl*_ and *c*_*mit*_ are related to the covariance between the fitness of a heterokaryon and the fitness of its acceptor homokaryon parent. For homokaryons, we define fitness as:

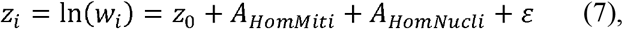

where *z*_*i*_ is the logarithm of the fitness of homokaryon *i*, *z*_0_ is the average homokaryon fitness, *A*_*HomMiti*_ is the homokaryon mitochondrial genetic value, *A*_*HomNucli*_ is the homokaryon nuclear genetic value and *ɛ* is the environmental error. Similarly, for heterokaryons, we define fitness as:

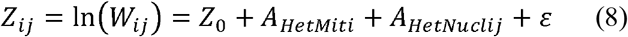

where *Z*_*ij*_ is the logarithm of the fitness of heterokaryons formed with parental homokaryons *i* and *j*, *Z*_0_ is the average heterokaryon fitness, *A*_*HetMiti*_ is the heterokaryon mitochondrial genetic value (only the acceptor *i* is providing the mitochondrion),*A*_*HetNuclij*_ is the heterokaryon nuclear genetic value and *ɛ* is the environmental error. We consider that heterokaryon mitochondrial and nuclear genetic effects are determined by a set of mutations that either have a fitness effect in homokaryons (hereafter *HomHet* superscript) or do not have a fitness effect in homokaryons (hereafter *HetOnly* superscript), such that:

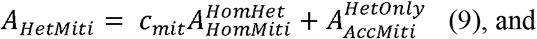

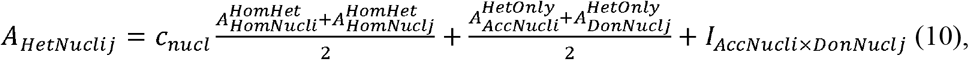

where *I*_*AccNucli*×*DonNucij*_ represents the acceptor x donor interaction and accounts for non-additive effects between acceptor and donor genomes (i.e. due to dominance and epistasic effects). This interaction can be decomposed as follows:

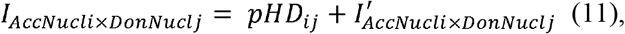

where 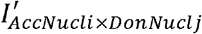 is the residual interaction after accounting for genetic distance (see analytical results above and Eq. (6) for the definition of *p*, *H* and *D*_*ij*_). After averaging measurements over multiple replicates (so that we can ignore environmental error), the slope of the regression of the heterokaryon value over the value of either its donor or acceptor homokaryon parent is:

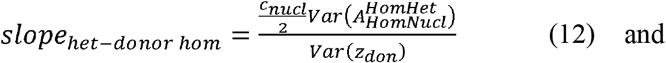

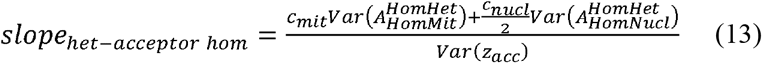

The variances, 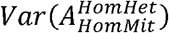 and 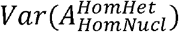 respectively represent the part of the variance among homokaryon genetic values determined by mitochondrial and nuclear loci that also have an effect in heterokaryons, whereas *Var*(*z*_*don*_) and *Var*(*z*_*acc*_) respectively represent the genetic variance among donor and acceptor homokaryons. When the fitness effects of nuclear and mitochondrial mutations between the two phases of the life cycle are independent, both *slope*_*het-donor hom*_ and *slope*_*het-acceptor hom*_ equal zero. When donor and acceptor homokaryons are randomly selected for heterokaryon synthesis, the variances between donor and acceptor homokaryons are likely to be identical (i.e. *Var*(*z*_*don*_) = *Var*(*z*_*acc*_)) and 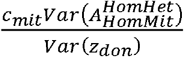 can be computed as *slope*_*het-acceptor hom*_ - *slope*_*het-donor hom*_. When there is no mitochondrial effects and no homokaryon-specific nuclear effects, 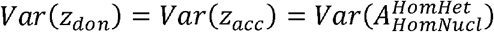, and the two regressions slopes equal 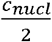 (see Supplemental Material File S1).

In diploid organisms, heterosis is usually referred to as a F1 performance that exceed the average parental performance (Lynch and Walsh 1998, p.222). By analogy, we define haploid mid-parent heterosis or heterokaryon vigor for fungi, *pH,* as the difference between a heterokaryon genetic effect (i.e. the sum of acceptor, donor genetic effects and their interaction) and the average of the parental homokaryon genetic effects after controlling for mean differences between homokaryons and heterokaryons (i.e. accounting for *c*_*nucl*_). Contrary to diploid midparent heterosis (defined as the difference between the genetic value of a diploid strain and the average genetic value of the two autozygous strains formed using its parents, e.g. Zörgö *et al.* 2012), haploid mid-parent heterosis (or heterokaryon vigor) can be estimated in self-incompatible species. Indeed, when there is no homokaryon-or heterokaryon-specific fitness effect, we have:

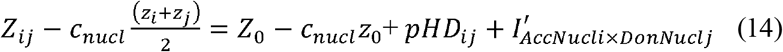

The slope of the increase in haploid mid-parent heterosis with pairwise genetic distance measured between the two homokaryon parents represents the parameter *pH* and is directly related to the average level of dominance (see analytical results above for a mathematical definition of *H*). We build graphs to illustrate the relationship between the MGR of a heterokaryon and the MGR of either its donor (Eq. 12) or acceptor (Eq. 13) homokaryon parent as well as the relationship between mid-parent heterosis (assuming that *c*_*nucl*_ =1, see result section below) and the genetic distance between parents (Eq. 14). To remove environmental effects, homokaryon and heterokaryon genetic effects were estimated by averaging the values measured across different replicates, resulting in 16 non-senescent homokaryon and 225 heterokaryon average MGR estimates. We also develop a custom statistical model to simultaneously estimate the coefficients *c*_*nucl*_, *c*_*mit*_ and *H* without any simplifying assumption (i.e. considering both homokaryon-and heterokaryon-specific nuclear and mitochondrial fitness effects) and fit the relationships predicted by Eq. 12, 13 and 14.

We implemented custom multiple-membership (Rode *et al.* 2017a) animal models (Henderson 1950; Lynch and Walsh 1998) using the *lme4* package (Bates *et al.* 2015) in R v3.4.4 (http://www.r-project.org/, R Development Core Team 2013). We used bivariate linear mixed models to partition genetic and environmental effects. Briefly, fixed effects comprised different intrinsic fitness effects for heterokaryons and homokaryons (*strain type* factor) and the genetic distance between parental homokaryons (set at zero for homokaryons, and at the genome-wide genetic distance between parental homokaryons for heterokaryons, *genetic distance* covariate). Random genetic effects where partitioned into genetic effects shared between a homokaryon and its heterokaryon offspring (i.e. to estimate *c*_*nucl*_ and *c*_*mit*_) and homokaryon-specific or heterokaryon specific genetic effects (see Supplemental File S1 for model details). For model selection, we used the corrected Akaike Information Criterion (AICc, Burnham and Anderson 2002). Models with AICc differences smaller than two compared to the model with lowest AICc (*ΔAICc* < 2) are considered as strongly supported by the data. Models with the same log-likelihood as the model with lowest AICc and that differed from it by a single variable were considered as not supported (Burnham and Anderson 2002 p.131). We used a three-step approach for model selection. First, we tested whether the covariance between nuclear (respectively mitochondrial) genetic effects is not null by selecting the model with the optimal structure for random effects. We compare six models with different combinations of covariance structure for nuclear and mitochondrial genetic effects (Table 2), but with the same fixed effects (including a genetic distance effect for *H*). As these six models comprise different random effect structures, we fit them using restricted maximum likelihood (REML, Zuur *et al.* 2009, p.121). Second, if the best model with in *ΔAICc* = 0 the first step includes *c*_*nucl*_ or *c*_*mit*_, we compare this best model with models where *c*_*nucl*_ or *c*_*mit*_ is set to one (i.e. we test if *c*_*nucl*_ ≠1 vs. *c*_*nucl*_ = 1 and if *c*_*mit*_ ≠ 1 vs. *c*_*mit*_ = 1, Table 3) using maximum likelihood (ML, Zuur *et al.* 2009, p.121). Third, we use the covariance structure of the best model in the first step and compare models including or lacking either the effect of genetic distance (i.e. test *H* ≠ 0 vs.*H* = 0) and/or including or lacking ploidy differences in mean fitness (i.e. test *mean homokaryons* ≠ *mean hetrokaryons* vs. *mean homokaryons* = *mean hetrokaryons*) using maximum likelihood (Table 4).

**Table 2.**
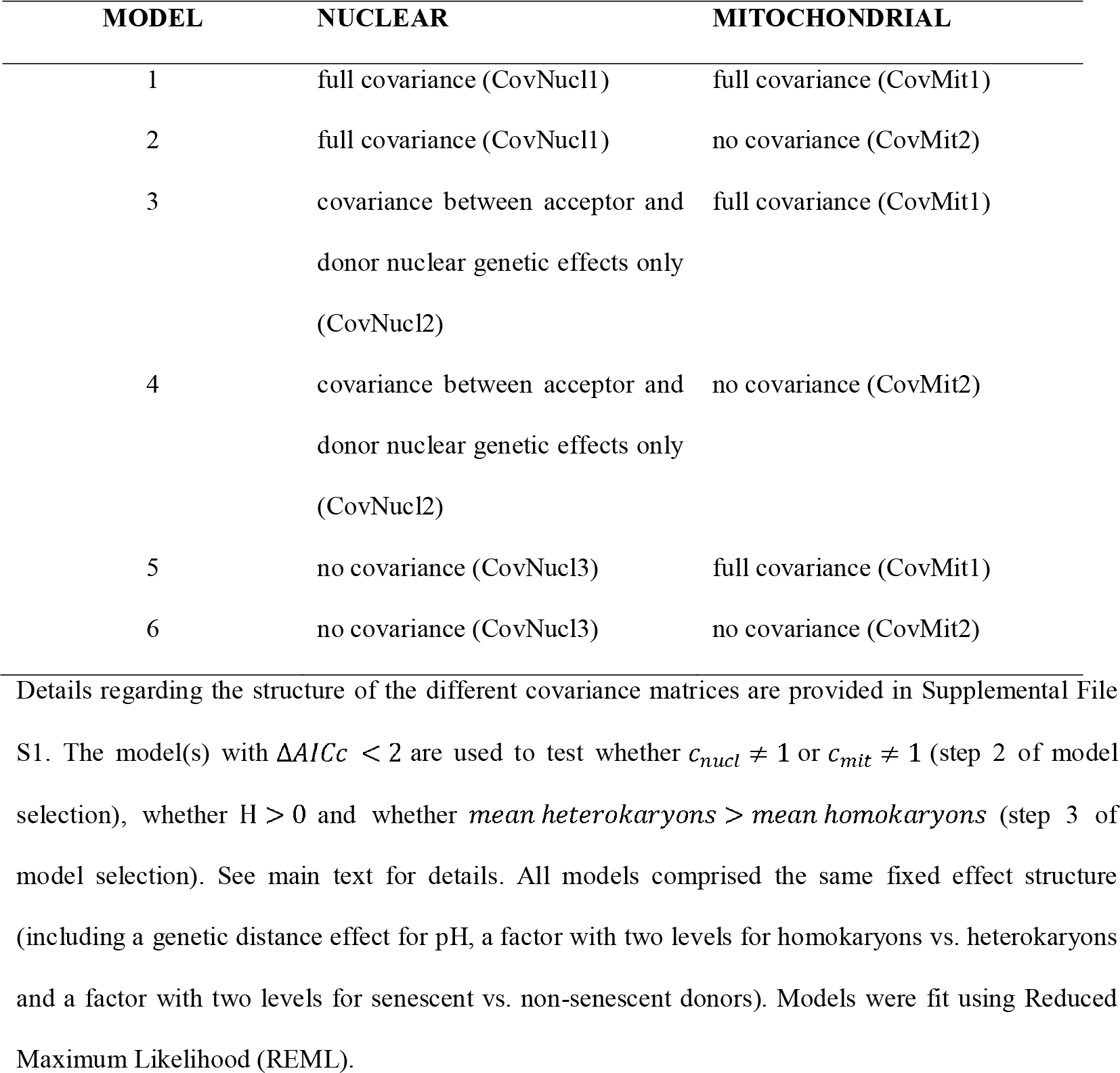
Covariance matrices of the six models with different structures for nuclear and mitochondrial genetic effects tested in step 1 of model selection.

**Table 3.** List of the different models to test whether *c*_*nucl*_ ≠ 1 and *c*_*mit*_ = 1 in step 2 of model selection when the best model in step 1 include (a) a covariance between homokaryon and acceptor or donor nuclear genetic effects or (b) a covariance between homokaryon and heterokaryon mitochondrial genetic effects.

|  | MODEL | EFFECT TESTED |
| --- | --- | --- |
| (a) | A1 | $c_{nucl} \neq 1$ |
| | A2 | $c_{nucl} = 1$ |
| (b) | B1 | $c_{mit} \neq 1$ |
| | B2 | $c_{mit} = 1$ |

Models were fit using Reduced Maximum Likelihood (REML).

**Table 4.**
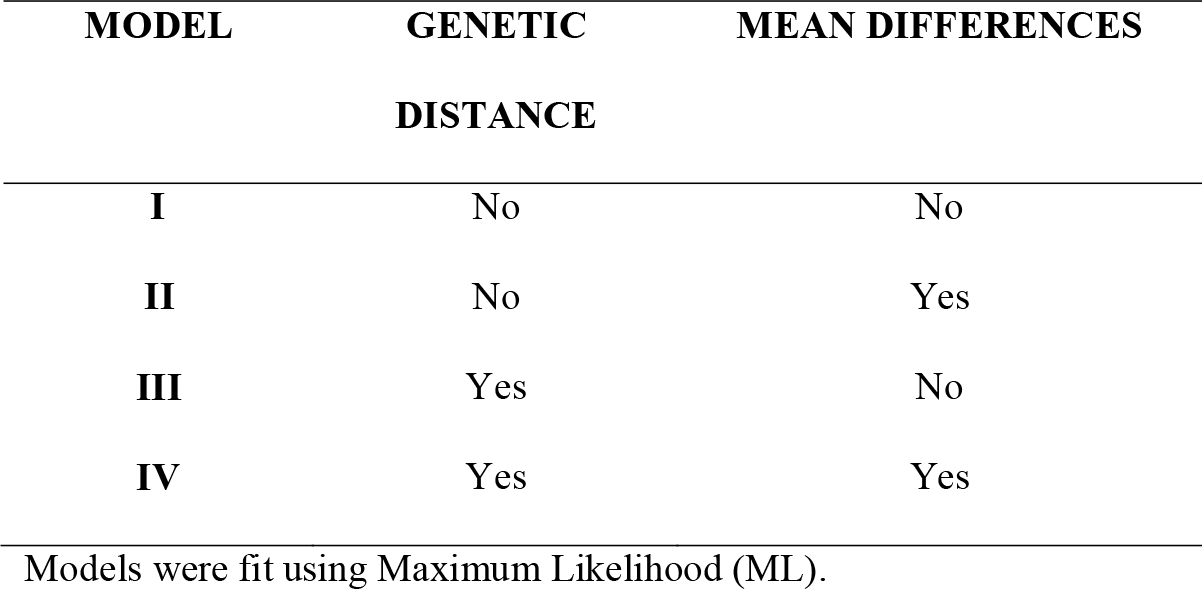
List of the four models to test for an effect of genetic distance and for mean differences between homokaryons and heterokaryons.

Parametric bootstrapping represents an alternative approach to AICc model selection, but performing the appropriate simulations for our full model would be very computationally intensive. We used this approach to compute the 95% confidence interval of the estimates of a reduced model selected based on AICc and of the proportion of homokaryon and heterokaryon phenotypic variance accounted by genetic (nuclear or mitochondrial) and environmental effects (percentile bootstrap method; Davison and Hinkley 1997; Bates *et al.* 2015). We also estimated our power of detecting that *c*_*nucl*_ differed from 1 using simulation-based power analyses based on our experimental design (Johnson *et al.* 2015). For both fixed effects and variance components, we used the estimates from the model with *c*_*nucl*_ ≠ 1 in step 2. We simulated 100 independent datasets for each value of *c*_*nucl*_ ranging from 0.1 to 2. We analysed each dataset using either a model where *c*_*nucl*_ was free to vary or a model where it was set to 1. Power was estimated as the proportion of the 100 datasets in which the model with *c*_*nucl*_ ≠ 1 was better than the other model (i.e. AIC(*model c*_*nucl*_ = 1) - AICc(model *c*_*nucl*_ ≠ 1) > 2. We used an arbitrary threshold power of 80% for the analyses (Johnson *et al.* 2015).

### Data availability

Supplemental Material Files S1-S5 available at Figshare (Clergeot & Rode *et al.* 2018): https://figshare.com/s/a19fa51f96a35c105684

File S1, Analytical and statistical models to estimate *c*_*nucl*_, *c*_*mit*_ and pH (Figures S1-S2). (.pdf)

File S2, Supplemental Material (Analysis of the ability to degrade wood) and Figures S3-S10 (.pdf)

File S3, Details regarding bioinformatics analyses. (.pdf)

File S4, Supplemental tables (.xlsx) with list of the isolates (Table S1), geographic distance matrix (Table S2), nuclear and mitochondrial genetic distance matrices (Tables S3-S4), AICc tables, best model estimates and proportion of variance explained for MGR (Table S5-S7) and the ability to degrade wood (WWL, Tables S8-S10).

File S5, ZIP archive with MGR and WWL phenotypic data and scripts needed to reproduce the results presented in this manuscript. In particular, the script to fit multiple membership statistical models developed for this study (MMmodel.Rmd). The reference genome, fastq files and VCF of the different isolate are deposited in the European Nucleotide Archive (ENA) Sequence Read Archive (SRA) (project accession PRJEB27090).

## Results

### Analysis of H. parviporum population structure

Based on STRUCTURE analyses, the best number of clusters was one. *Fst* values between pairs of populations were consistently low and non-significant (Table 5). Hence, we did not detect any population structure among our homokaryon isolates. This finding is in line with the absence of population structure found in the related species *H. annosum* s.s. (Dalman *et al.* 2013).

**Table 5.** Pairwise non-significant *Fst* values between the different geographic areas.

|  | **GEOGRAPHIC**<br>**AREA 1** | **GEOGRAPHIC**<br>**AREA 2** | **GEOGRAPHIC**<br>**AREA 3** | **GEOGRAPHIC**<br>**AREA 4** |
| --- | --- | --- | --- | --- |
| **GEOGRAPHIC**<br>**AREA 1** |  | 0.006 | 0.019 | 0.027 |
| **GEOGRAPHIC**<br>**AREA 2** | 0.006 |  | 0.032 | 0.015 |
| **GEOGRAPHIC**<br>**AREA 3** | 0.019 | 0.032 |  | 0.056 |
| **GEOGRAPHIC**<br>**AREA 4** | 0.027 | 0.015 | 0.056 |  |

See Table S1 for the list of isolates per geographic area.

We found 11 different mitochondrial haplotypes among our 16 non-senescent homokaryon isolates (senescent homokaryons that never behaved as acceptor were not considered for the analyses of mitochondrial genetic distance). Most mitochondrial haplotypes were unique to a single sampling area (except for two haplotypes, Table S1). Genetic distance between pairs of mitochondrial haplotypes was partially positively correlated with nuclear genetic distance between pairs of homokaryons carrying these haplotypes (Mantel-test, *R*=0.43, *P*=0.01, Figure S5).

### Average effect of ploidy and average level of dominance

Results regarding the ability to degrade wood, used a second fitness-related trait, are presented in Supplemental File S2. The MGR of heterokaryon isolates (mean=6.3 mm/day, sd^2^=1.4 mm/day) was higher on average and less variable than the MGR of homokaryon isolates (mean=5.3 mm/day, sd^2^=4.1 mm/day, Figure 3).

**Figure 3.**
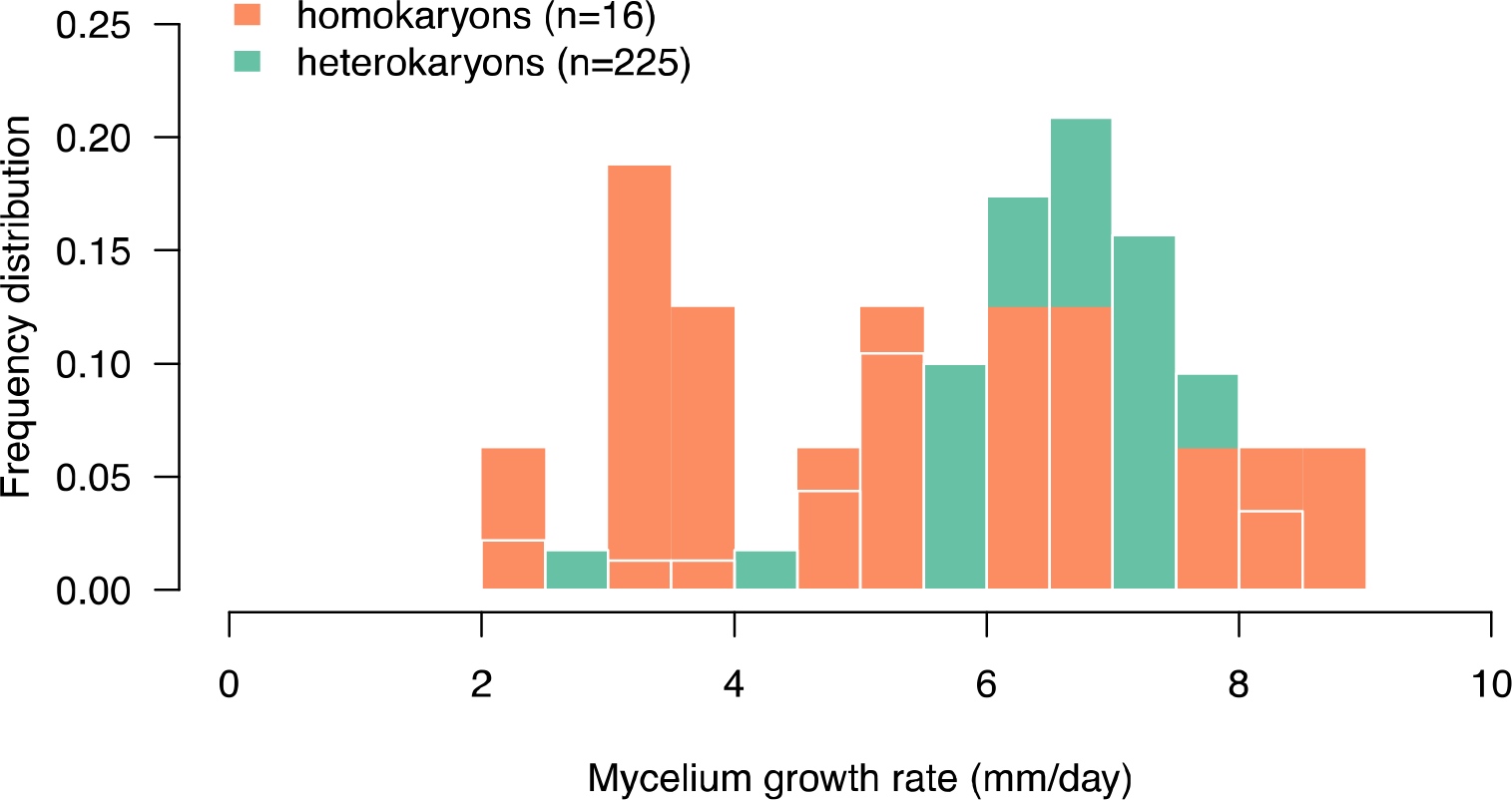
Distribution of mycelium growth rate for homokaryon and heterokaryon isolates.

The MGR of heterokaryons increased with the MGR of their donor or acceptor homokaryon parent (Figure 4). Consistent with this observation, models 1 and 2 that included a covariance between homokaryon, acceptor and donor genetic effects had high support (*ΔAlCc* > 2 for other models, Table S5A). The estimate of *c*_*nucl*_ was lower than one (Table S6, ML estimate for *c*_*nucl*_ =0.70, 95% confidence interval = 0.37, 1.15), but with low support (*ΔAlCc* = 0.60 and 0 for models A1 and A2 with *c*_*nucl*_ = 0.69 and 1 respectively, Table S5B). Retrospective power analyses showed that our statistical power to detect whether *c*_*nucl*_ differed from one was relatively low provided our experimental design (power was below 80% for values of *c*_*nucl*_ below 1.7, Figure S8). Overall, the proportion of phenotypic variance explained by acceptor and donor nuclear genetic effects was lower than the one explained by homokaryon genetic effect (32% vs. 46%, Table S7).

**Figure 4.**
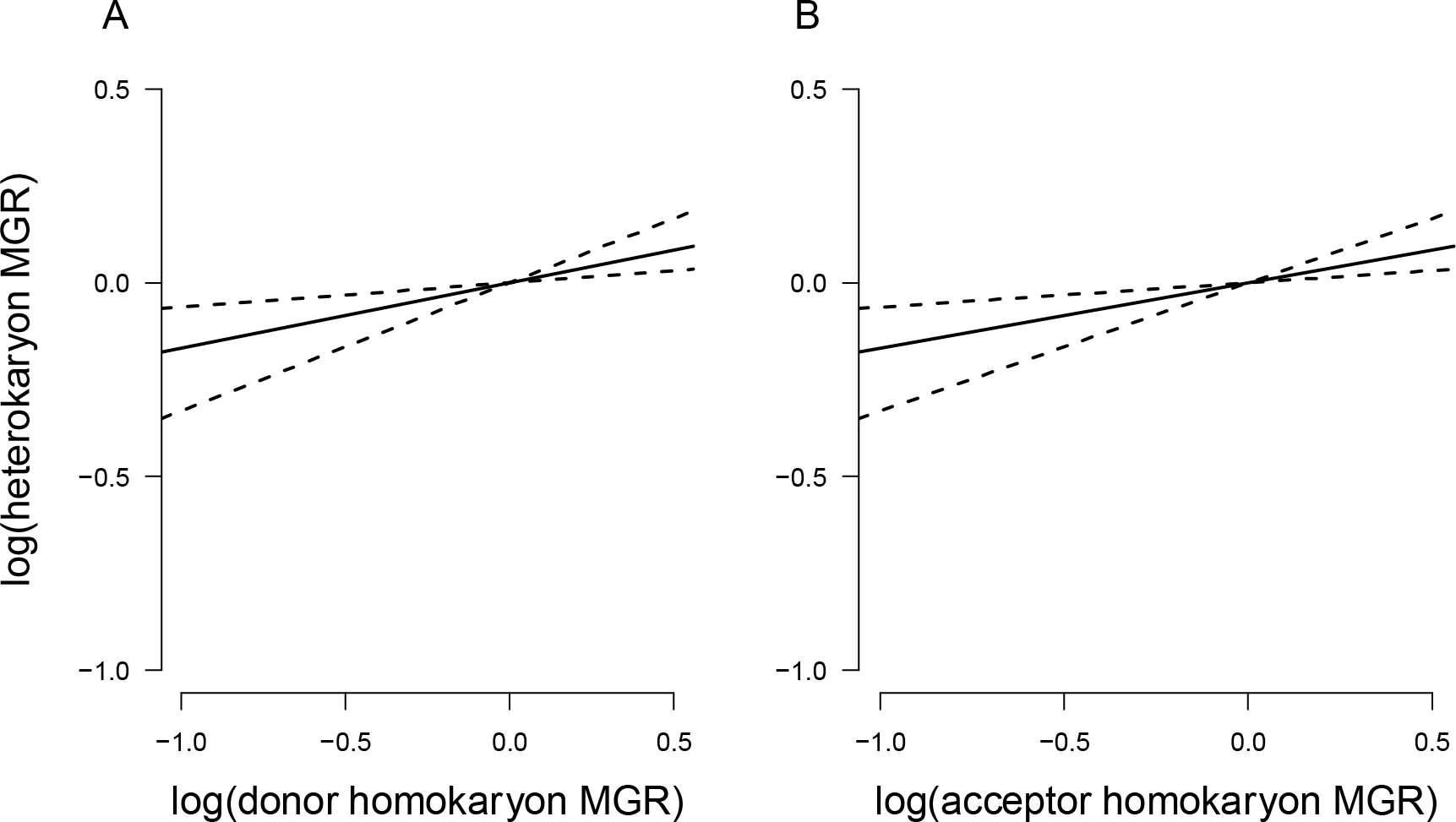
Relationship between the genetic values of heterokaryon offspring and the genetic values of either (A) their donor or (B) their acceptor homokaryon parent. As the covariance between homokaryon and heterokaryon mitochondrial genetic effects was zero, the slopes in A and B are identical. Solids lines represent the fitted slope between heterokaryons and donor or acceptor homokaryon parent. Dashed lines represent the 95% confidence interval of the slope estimate based on 100 parametric bootstrap values. Genetic values are represented on a log scale and mean-centered (n=171 or n=224 heterokaryons with donor or acceptor homokaryon MGR data respectively).

Model 1, that included a covariance between mitochondrial genetic effects in homokaryons and heterokaryons 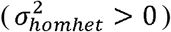, had *ΔAlCc* = 1.81 compared to model 2 without this covariance (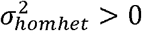, *ΔAlCc* = 0, Table S5A), suggesting that covariance between homokaryon and heterokaryon mitochondrial genetic effects had low support. The log-likelihoods of the models that included an effect of genetic distance and/or and different average MGRs for homokaryons and heterokaryons was the same has the model without these effects (Table S5C, Figure 5), so this effect had no support. Mitochondrial effects accounted for 35% of phenotypic variation in MGR among homokaryons, but only 3% of phenotypic variation among heterokaryons (Table S7).

**Figure 5.**
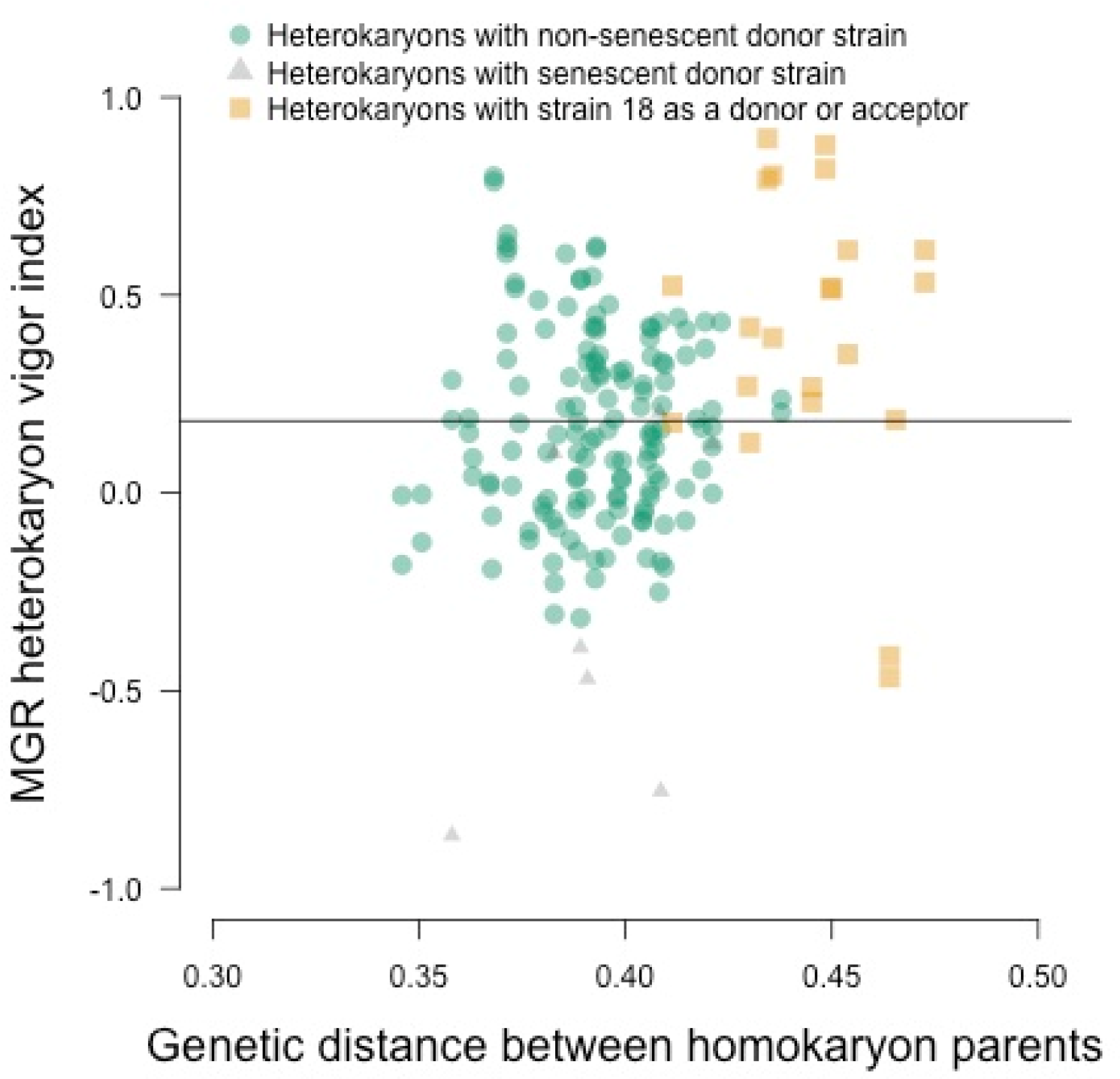
Relationship between MGR heterokaryon vigor index and genetic distance between homokaryon. Strain 18 is more genetically distant compared to the other strains and heterokaryon synthetized using this strain have higher trait values on average, creating a spurious correlation between heterokaryon and genetic distance when the non-independence between heterokaryons is not accounted for. Heterokaryon vigor is computed as the difference between log (heterokaryons value) and log (homokaryon mid-parent value). Solid lines represent the average heterokaryon vigor value (n=175 estimates).

## DISCUSSION

### A new statistical framework to estimate fitness effects across ploidy phases of the life cycle

Recent theoretical models predict that in haploid-diploid species, intrinsic fitness during the diploid phase should be higher than during the haploid phase, and that the average of the ratio of the fitness effect in homozygous diploids over fitness effect in haploids across mutations should be higher than one (i.e. c_nucl_ > 1, assuming mutations are recessive on average; Scott and Rescan 2017). We derived a new statistical model to jointly test these predictions based on the covariance between the mean fitness of the haploid parents (e.g. homokaryon parents in fungi) and the fitness of their diploid offspring (e.g. heterokaryons in fungi). Our model accounts for important factors that could potentially bias the estimation of these genetic parameters (e.g. phenotypic variation due to nuclear or mitochondrial mutations with haploid-or diploid-specific fitness effects or the senescent phenotype of some strains).

Our statistical framework presents several advantages compared to previous methods. First, through the simultaneous estimation of the average level of dominance and of *c*_*nucl*_, it can be used to directly test for intrinsic fitness differences between diploid and haploid individuals, while controlling for the masking of deleterious mutations in diploids. Failing to account for such confounding effect have been a recurrent issue in many studies investigating fitness differences between haploids and diploids in self-incompatible species (Scott and Rescan 2017). As a consequence, the most accurate quantification of ploidy effects on fitness have been restricted to self-compatible species such as yeast (e.g. Korona 1999; Zörgö *et al.* 2013). Our case study in *H. parviporum* shows that an accurate quantification of these effects in self - incompatible species is possible. Second, our quantitative genetic framework controls for mitochondrial genetic effects and acceptor x donor interactions, and accounts for the non-independence between diploid hybrids that share one haploid parent. These confounding factors could strongly bias estimates based on conventional methods (e.g. Korona 1999; Zörgö *et al.* 2012; Plech *et al.* 2014; Shapira *et al.* 2014; Stelkens *et al.* 2014; Bernardes *et al.* 2017). One isolate (strain 18) was on average more distantly related to the other isolates and had higher than average donor and acceptor genetic effects, creating a spurious correlation between heterokaryon vigor and genetic distance (Figure 4). By accounting for the non-independence between heterokaryons formed with this strain, our model showed that genetic distance between homokaryon parents did not affect heterokaryon MGR. Third, our model clearly separates acceptor nuclear effects from mitochondrial effects for loci that are expressed both in homokaryons and heterokaryons, provided that some reciprocal crosses are successful (i.e. that some heterokaryons or diploids have the same nuclear background but different mitochondrial haplotypes). Our model can also separate nuclear and mitochondrial genetic effects that are either homokaryon-or heterokaryon-specific, provided that the correlation between matrices of mitochondrial and nuclear genetic distances is low, i.e. that some haploids (or homokaryons) have similar mitochondrial haplotypes but different nuclear backgrounds and vice versa. Fourth, by using a quantitative genetic framework, our model can refine the understanding of the genetic architecture of fitness in haploids and in diploids. Indeed, some genes could affect only haploid fitness, only diploid fitness, or both (Rescan *et al.* 2016), resulting in differences in mean fitness between haploids and diploids, or in differences in the variance in fitness observed among haploids and among diploids. For example, the expression of a gene only at one stage could result in intrinsic fitness differences between diploids and haploids, while a different transcription pattern between stages could result in a difference in the variance in fitness.

Our model could potentially suffer from the same limitations as methods used to measure inbreeding depression or to infer relatedness using molecular markers (e.g. Szulkin *et al.* 2010). It assumes that genome-wide genetic distance is a good approximation of the genetic distance between selected loci of the two haploid parents. This assumption could be wrong whenever mutations that affect fitness are non-evenly distributed across the genome. Whenever mutations cluster together in some specific genomic regions, genetic distance should be estimated in those genomic regions only. Using programs to classify mutations as neutral or deleterious (Adzhubei *et al.* 2010; Vaser *et al.* 2016) could potentially alleviate this limitation. A second possible limitation is that our main statistical model does not account for epistatic interactions between mutations. In order to investigate this issue, we derived a more complex model that accounts for pairwise epistatic interactions (see Supplemental Material File S1). This model shows that Eq. 5 and 6 are accurate when the mean of epistatic effects is zero. Such epistasis pattern is observed in some experimental data and is expected under Fisher’s Geometric model of adaptation (Martin *et al.* 2007). However, when epistasis is antagonistic, our model underestimates *c*_*nucl*_ and overestimates *H,* while the reverse is true when epistasis is synergistic (Supplemental Material File S1). Using F2 crosses and the appropriate statistical extension would allow estimating epistatic effects. Our model and our proof of principle simulations could be used as starting points for such developments.

### Investigations of theoretical predictions in natural isolates of *H. parviporum*

In basidiomycete fungi, intrinsic fitness differences can arise due to differences between homokaryons and heterokaryons in physiology (e.g. formation of clamp connections in heterokaryons, Simchen and Jinks 1964), in the number of nuclei or mitochondria per cell (Hansen 1979) or in gene expression (Meng *et al.* 2013; Liu *et al.* 2017). Theoretical models predict that intrinsic fitness should be slightly higher in diploids than in haploids in haploid-diploid species (Scott and Rescan 2017). Despite a higher MGR in heterokaryons compared to homokaryons (Figure 3), we failed to find significant intrinsic fitness differences between these two stages. As theory predicts these differences to be small, detecting them experimentally might prove challenging.

We fail to find support for an effect of genetic distance on MGR (i.e. models where *H* differed from zero had no support). Two alternative explanations could explain this pattern. First, deleterious mutations that segregate in natural populations of *H. parviporum* might be co-dominant on average in heterokaryons (i.e. *H* = 0 and *h* = 0.5). This observation might appear at odds with theoretical and experimental studies showing that random mutations are partially recessive (h~0.27; Manna *et al.* 2011). However, mildly deleterious mutations are segregating at higher frequencies than strongly deleterious mutations in natural populations (Scott and Rescan 2017). Hence, if these mildly deleterious mutations are less recessive than strongly deleterious ones (as often observed in experimental studies; e.g. Marek and Korona 2016), we expect the average level of dominance of segregating mutations to be closer to 0.5 (even when random mutations have h=0.27). This hypothesis could be tested using our model to estimate H using natural isolates (i.e. for segregating mutations) or mutation accumulations lines (i.e. for random mutations). Second, deleterious mutations could be partially recessive in heterokaryons, but their masking could only be visible when crossing closely-related parents. Indeed, it is possible that heterokaryon fitness increases with genetic distance and plateaus after a threshold. As our crosses between the six most closely related homokaryons failed (i.e. crosses with genetic distance lower than 0.34), we had low power to detect such a saturating effect. In agreement with this hypothesis, previous studies in yeast have found no effect of genetic distances on the fitness of intraspecific hybrids from crosses between wild isolates (Zörgö *et al.* 2012; Shapira *et al.* 2014; Bernardes *et al.* 2017), but a positive effect for hybrids from crosses between more closely related domesticated isolates (Plech *et al.* 2014). In addition, potential epistatic effects could bias downward the estimates of *H* (see above). Testing additional heterokaryons, synthetized using more closely related parents, should help discriminate between these alternative hypotheses.

Contrary to theoretical expectations, we found that *c*_*nucl*_ was not higher than one. Although our findings of *c*_*nucl*_ = 1 and 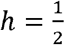 is consistent theoretical predictions based on the observation that the heterokaryon phase is the longest in this species (Johannesson and Stenlid 2004), we cannot rule out that a lack of statistical power is responsible for these findings. Our retrospective power analyses showed that we could only detect values of *c*_*nucl*_ larger than 1.7. Increasing our statistical power using more homokaryons isolates would help verify these results. For the ability to degrade wood, we had fewer replicates than for MGR which might explain the lack of covariance between heterokaryon and acceptor and donor homokaryon parents (Supplemental File S2, Figure S9).

### Genetic architecture of fitness traits in *H. parviporum*

Our results also provide important insights into the genetic architecture of our two fitness-related traits in *H. parviporum.* The genetic architecture we unravelled for MGR appears similar to the genetic architecture of growth rate in yeast. Indeed, the variance in fitness is higher among haploids than among diploids in *S. cerevisiae* (Zörgö *et al.* 2012), and diploid fitness is highly correlated to haploid fitness (Korona 1999; Zörgö *et al.* 2013). However, these studies did not investigate relative contribution of nuclear vs. mitochondrial effects to haploid and diploid phenotypic variance.

Our most striking result is that both mitochondrial and nuclear genetic variation is much greater among homokaryons than among heterokaryons. If *c*_*nucl*_ = 1, we expect nuclear genetic variation among heterokaryons to be half of nuclear genetic variation among homokaryons. However, genetic variation was an order of magnitude lower among heterokaryons than among homokaryons (Table S6). In addition, genetic variation among mitochondrial haplotypes was also larger among homokaryons than heterokaryons (Table S6). To our knowledge, this pattern has not been documented in previous basidiomycete studies that relied on a large number of highly related homokaryons and heterokaryons (i.e. derived from one or few crosses, Simchen and Jinks 1964; Williams *et al.* 1976; Elliott *et al.* 1979). Two non-mutually exclusive hypotheses could explain these observations. First, we expect the variance among homokaryons to be larger than the variance among heterokaryons when *c*_*nucl*_ < 1 or *c*_*mit*_ < 1. Second, the genetic architecture of fitness might differ between homokaryons and heterokaryons (e.g. if the effects of mutations in homokaryons and heterokaryons are not correlated). The second hypothesis appears more likely, as the model with lowest AICc had *c*_*nucl*_ = 1 and did not include any covariance between mitochondrial effects in heterokaryons and parental homokaryons. We acknowledge that mitochondrial and nuclear effects with specific effects in homokaryons or in acceptors are difficult to disentangle in our design as nuclear and mitochondrial genetic distance matrices were positively correlated (R=0.43) and as most mitochondrial haplotypes were carried by a single homokaryon. However, this lack of identifiability does not comprise our observation that genetic variances are much larger among homokaryons than among heterokaryons.

Previous studies in basidiomycetes (e.g. James *et al.* 2008) found phenotypic differences between heterokaryons having the same nuclear background but different cytoplasms. Our model helps to better understand the different factors potentially responsible for these differences. First, these differences could be due to mitochondrial effects, as they account for 3% of phenotypic variance in MGR in our experiment. Second, different genetic architecture for acceptor and donor genetic effects could potentially explain such differences. For example, the variance of nuclear genetic effects is larger among donors than among acceptors for the ability to degrade wood (Table S10). Third, such differences could be due to cytoplasmic factors that are differently transmitted by donors and acceptors. Heterokaryon syntheses using homokaryons with a senescent phenotype failed most of the time when the homokaryon was used as an acceptor, but not when it was used as a donor. We also found that homokaryons with a senescent phenotype produce heterokaryons that are less fit on average. Although they grew normally in 2012/2013, the senescent phenotype was observed in some or all of the precultures of 18 isolates in 2015-2018. The probability of a preculture harboring this senescence phenotype increases with the age of a colony (Stenlid and Rayner 1989). These observations suggest that senescence does not represent a direct nuclear effect, and that some epigenetic effects are transmitted by donor nuclei, or that some additional genetic materials beside donor nuclei (e.g. prions or viruses) are acquired during the storage and are transferred to heterokaryons. Importantly, senescence effects did not bias our estimates of *c*_*nucl*_ and *c*_*mit*_, as the covariance between heterokaryon and their homokaryon parents is only estimated using non-senescent homokaryons. Fourth, these differences could arise, for a given isolate, when the average proportion of nuclei within the heterokaryon mycelium changes depending on whether it is used as a donor or as an acceptor in a cross. This is however unlikely here, as a previous study in *H. parviporum* showed that the proportions of nuclei within a heterokaryon when a given isolate is used as a donor or as an acceptor are highly correlated (James *et al.* 2008). For other species where the proportion of acceptor and donor nuclei might not be correlated, pool-sequencing can be used to reliably estimate the frequency of each nucleus (Rode *et al.* 2017b) and our model could easily be modified to take different acceptor and donor nuclear ratios into account.

### Implication for the evolution of life cycles

A promising line of research would be to use our model to test whether effective dominance is lower in species where the haploid phase of the life cycle is predominant (e.g. *Aspergillus* spp., *Neurospora* spp., *Schizosaccharomyces* spp., etc.) compared to species where the diploid phase is predominant (e.g. *Saccharomyces cerevisiae, Schizophyllum commune,* etc.). Indeed, everything else being equal, we expect the proportion of the life cycle spent as diploid to correlate positively with *hc_nuc_i.* Hence, using our model to compare these parameters among species that differ in the proportion of their life cycle spent as diploid, represents a promising area of research. Importantly, selfing, clonality, and ecological differences between haploids and diploids represent factors that could also explain the maintenance of haploid-diploid life cycles beside effective dominance (Rescan *et al.* 2016; Scott and Rescan 2017). Hence, our model could help better understand the relative importance of these factors in maintaining haploid-diploid life cycles.

## Acknowledgements

We thank Marie Rescan for very helpful discussions on theoretical models regarding the evolution of haploid-diploid life cycles and Jean-Nicolas Jasmin for suggestions regarding yeast literature. We are grateful to Aneil Agrawal and two anonymous reviewers whose comments helped improve previous versions of the manuscript. This work was supported by the Swedish Foundation for Strategic Research, grant no. RBb08-0011 and Carl Trygger Foundation (PHC, ÅO, MBD) as well as the CeMEB LabEx/University of Montpellier, grant no. ANR-10-LABX-04-01 (NOR).

## Authors contributions

SG and NOR developed and implemented the analytical quantitative genetic model. NOR performed the simulations and the statistical analyses. KI and MBD performed DNA extractions and genome sequencing. PHC and MBD performed the bioinformatics analyses. ÅO and PHC planned the experiment and performed homokaryon crosses, mycelium growth rate and wood weight loss assays. NOR wrote the manuscript with inputs from the coauthors.

## Literature cited

Adzhubei I. A., Schmidt S., Peshkin L., Ramensky V. E., Gerasimova A., et al, 2010 A method and server for predicting damaging missense mutations. Nat. Methods 7: 248.

Bates D., Mächler M., Bolker B., Walker S., 2015 Fitting Linear Mixed-Effects Models Using lme4. J. Stat. Softw. 67: 1–48.

Beadle G. W., Coonradt V. L., 1944 Heterocaryosis in Neurospora crassa. Genetics 29: 291–308.

Bernardes J. P., Stelkens R. B., Greig D., 2017 Heterosis in hybrids within and between yeast species. J. Evol. Biol. 30: 538–548.

Burnham K. P., Anderson D. R., 2002 Model selection and multimodel inference: a practical information-theoretic approach. Springer Science & Business Media, New York, NY.

Chin C.-S., Alexander D. H., Marks P., Klammer A. A., Drake J., et al., 2013 Nonhybrid, finished microbial genome assemblies from long-read SMRT sequencing data. Nat. Methods 10: 563.

Chippindale A. K., Gibson J. R., Rice W. R., 2001 Negative genetic correlation for adult fitness between sexes reveals ontogenetic conflict in Drosophila. Proc.Natl. Acad. Sci. 98: 1671–1675.

Coelho S. M., Peters A. F., Charrier B., Roze D., Destombe C., et al., 2007 Complex life cycles of multicellular eukaryotes: new approaches based on the use of model organisms. Gene 406: 152–170.

Crockatt M. E., Pierce G. I., Camden R. A., Newell P. M., Boddy L., 2008 Homokaryons are more combative than heterokaryons of Hericium coralloides. Fungal Ecol. 1: 40–48.

Dalman K., Himmelstrand K., Olson Å., Lind M., Brandström-Durling M., et al, 2013 A genome-wide association study identifies genomic regions for virulence in the non-model organism Heterobasidion annosum ss. PLoS One 8: e53525.

Dassow P. Von, Ogata H., Probert I., Wincker P., Silva C. Da, et al., 2009 Transcriptome analysis of functional differentiation between haploid and diploid cells of Emiliania huxleyi, a globally significant photosynthetic calcifying cell. Genome Biol. 10: R114.

Davison A. C., Hinkley D. V., 1997 Bootstrap methods and their application. Cambridge university press, Cambridge.

Day P. R., Roberts C. F., 1969 Complementation in dikaryons and diploids of Coprinus lagopus. Genetics 62: 265–270.

Elliott C. G., Abou-Heilah A. N., Leake D. L., Hutchinson S. A., 1979 Analysis of wood-decaying ability of monokaryons and dikaryons of Serpula lacrymans. Trans. Br. Mycol. Soc. 73: 127–133.

Garbelotto M., Lee H., Slaughter G., Popenuck T., Cobb F. W., et al., 1997 Heterokaryosis is not required for virulence of Heterobasidion annosum. Mycologia 89: 92–102.

Garrison E., Marth G., 2012 Haplotype-based variant detection from short-read sequencing. arXiv Prepr. arXiv1207.3907.

Gerstein A. C., 2013 Mutational effects depend on ploidy level: all else is not equal. Biol. Lett. 9: 20120614.

Hansen E. M., 1979 Nuclear condition and vegetative characteristics of homokaryotic and heterokaryotic isolates of Phellinus weirü. Can. J. Bot. 57: 1579–1582.

Henderson C. R., 1950 Estimation of genetic parameters. Ann. Math. Stat. 21: 309–310.

Ihrmark K., Johannesson H., Stenström E., Stenlid J., 2002 Transmission of doublestranded RNA in Heterobasidion annosum. Fungal Genet. Biol. 36: 147–154.

James T. Y., Stenlid J., Olson Å., Johannesson H., 2008 Evolutionary significance of imbalanced nuclear ratios within heterokaryons of the basidiomycete fungus Heterobasidion parviporum. Evolution (N. Y). 62: 2279–2296.

Johannesson H., Stenlid J., 2004 Nuclear reassortment between vegetative mycelia in natural populations of the basidiomycete Heterobasidion annosum. Fungal Genet. Biol. 41: 563–570.

Johnson P. C. D., Barry S. J. E., Ferguson H. M., Müller P., 2015 Power analysis for generalized linear mixed models in ecology and evolution. Methods Ecol. Evol. 6: 133–142.

Korona R., 1999 Unpredictable fitness transitions between haploid and diploid strains of the genetically loaded yeast Saccharomyces cerevisiae. Genetics 151: 77–85.

Langmead B., Salzberg S. L., 2012 Fast gapped-read alignment with Bowtie 2. Nat Meth 9: 357–359.

Lind M., Nest M. van der, Olson Å., Brandström-Durling M., Stenlid J., 2012 A 2nd generation linkage map of Heterobasidion annosum sl based on in silico anchoring of AFLP markers. PLoS One 7: e48347.

Liu T., Li H., Ding Y., Qi Y., Gao Y., et al, 2017 Genome-wide gene expression patterns in dikaryon of the basidiomycete fungus Pleurotus ostreatus. brazilian J. Microbiol. 48: 380–390.

Lynch M., Walsh B., 1998 Genetics and analysis of quantitative traits. Sinauer Sunderland, MA.

Mable B. K., 2001 Ploidy evolution in the yeast Saccharomyces cerevisiae: a test of the nutrient limitation hypothesis. J. Evol. Biol. 14: 157–170.

Manna F., Martin G., Lenormand T., 2011 Fitness landscapes: an alternative theory for the dominance of mutation. Genetics 189: 923–937.

Marek A., Korona R., 2016 Strong dominance of functional alleles over gene deletions in both intensely growing and deeply starved yeast cells. J. Evol. Biol.

Martin G., Elena S. F., Lenormand T., 2007 Distributions of epistasis in microbes fit predictions from a fitness landscape model. Nat. Genet. 39: 555.

McBride R., Duncan G., Michael T., 2008 Fungal viral mutualism moderated by ploidy. Evolution (N. Y). 62: 2372–2380.

Meng L., Yan J., Xie B., Li Y., Chen B., et al, 2013 Genes encoding FAD-binding proteins in Volvariella volvacea exhibit differential expression in homokaryons and heterokaryons. Microbiol. Res. 168: 533–546.

Olson Å., Aerts A., Asiegbu F., Belbahri L., Bouzid O., et al., 2012 Insight into trade-off between wood decay and parasitism from the genome of a fungal forest pathogen. New Phytol. 194: 1001–1013.

Plech M., Visser J. A. G. M. de, Korona R., 2014 Heterosis is prevalent among domesticated but not wild strains of Saccharomyces cerevisiae. G3 Genes, Genomes, Genet. 4: 315–323.

Pringle A., Taylor J., 2002 The fitness of filamentous fungi. Trends Microbiol. 10: 474–481.

R Development Core Team R., 2013 R: A Language and Environment for Statistical Computing (RDC Team, Ed.). R Found. Stat. Comput. Vienna, Austria.

Raper J. R., 1966 Genetics of sexuality in higher fungi Ronald Press. New York, USA.

Redfern B., Pratt E., Gregory C., MacAskill A., 2001 Natural infection of Sitka spruce thinning stumps in Britain by spores of Heterobasidion annosum and long-term survival of the fungus. Forestry 74: 53–71.

Rescan M., Lenormand T., Roze D., 2016 Interactions between genetic and ecological effects on the evolution of life cycles. Am. Nat. 187: 19–34.

Rode N. O., Soroye P., Kassen R., Rundle H. D., 2017a Air-borne genotype by genotype indirect genetic effects are substantial in the filamentous fungus Aspergillus nidulans. Heredity (Edinb). 119: 1–7.

Rode N. O., Holtz Y., Loridon K., Santoni S., Ronfort J., et al., 2017b How to optimize the precision of allele and haplotype frequency estimates using pooled-sequencing data. Mol. Ecol. Resour. 18: 194–203.

Rokitta S. D., Nooijer L. J. de, Trimborn S., Vargas C. de, Rost B., et al., 2011 Transcriptome analyses reveal differential gene expression patterns between the life-cycle stages of Emiliania huxleyi (Haptophyta) and reflect specialization to different ecological niches. J. Phycol. 47: 829–838.

Schielzeth H., 2010 Simple means to improve the interpretability of regression coefficients. Methods Ecol. Evol. 1: 103–113.

Scott M. F., Rescan M., 2017 Evolution of haploid-diploid life cycles when haploid and diploid fitnesses are not equal. Evolution (N. Y). 71: 215–226.

Shapira R., Levy T., Shaked S., Fridman E., David L., 2014 Extensive heterosis in growth of yeast hybrids is explained by a combination of genetic models. Heredity (Edinb). 113: 316.

Simchen G., Jinks J. L., 1964 The determination of dikaryotic growth rate in the basidiomycete Schizophyllum commune: a biometrical analysis. Heredity (Edinb). 19: 629.

Stelkens R. B., Brockhurst M. A., Hurst G. D. D., Miller E. L., Greig D., 2014 The effect of hybrid transgression on environmental tolerance in experimental yeast crosses. J. Evol. Biol. 27: 2507–2519.

Stenlid J., 1985 Population structure of Heterobasidion annosum as determined by somatic incompatibility, sexual incompatibility, and isoenzyme patterns. Can. J. Bot. 63: 2268–2273.

Stenlid J., Rayner A. D. M., 1989 Tansley Review No. 19 Environmental and endogenous controls of developmental pathways: variation and its significance in the forest pathogen, Heterobasidion annosum. New Phytol. 113: 245–258.

Szafraniec K., Wloch D. M., Sliwa P., Borts R. H., Korona R., 2003 Small fitness effects and weak genetic interactions between deleterious mutations in heterozygous loci of the yeast Saccharomyces cerevisiae. Genet. Res. (Camb). 82: 19–31.

Szulkin M., Bierne N., David P., 2010 Heterozygosity-fitness correlations: a time for reappraisal. Evolution (N. Y). 64: 1202–1217.

Thornber C. S., 2006 Functional properties of the isomorphic biphasic algal life cycle. Integr. Comp. Biol. 46: 605–614.

Vaser R., Adusumalli S., Leng S. N., Sikic M., Ng P. C., 2016 SIFT missense predictions for genomes. Nat. Protoc. 11: 1.

Williams S., Verma M. M., Jinks J. L., Brasier C. M., 1976 Variation in a natural population of Schizophyllum commune. Heredity (Edinb). 37: 365.

Xu J., 1995 Analysis of inbreeding depression in Agaricus bisporus. Genetics 141: 137–145.

Zörgö E., Gjuvsland A., Cubillos F. A., Louis E. J., Liti G., et al, 2012 Life history shapes trait heredity by accumulation of loss-of-function alleles in yeast. Mol. Biol. Evol. 29: 1781–1789.

Zörgö E., Chwialkowska K., Gjuvsland A. B., Garré E., Sunnerhagen P., et al., 2013 Ancient evolutionary trade-offs between yeast ploidy states. PLoS Genet. 9: e1003388.

Zuur A., Ieno E. N., Walker N., Saveliev A. A., Smith G. M., 2009 Mixed effects models and extensions in ecology with R Springer Science & Business Media, New York, NY.

